# Accelerometry data in health research: challenges and opportunities: Review and examples

**DOI:** 10.1101/276154

**Authors:** Marta Karas, Jiawei Bai, Marcin Strączkiewicz, Jaroslaw Harezlak, Nancy W. Glynn, Tamara Harris, Vadim Zipunnikov, Ciprian Crainiceanu, Jacek K. Urbanek

## Abstract

Wearable accelerometers provide detailed, objective, and continuous measurements of physical activity (PA). Recent advances in technology and the decreasing cost of wearable devices led to an explosion in the popularity of wearable technology in health research. An ever increasing number of studies collect high-throughput, sub-second level raw acceleration data. In this paper we discuss problems related to the collection and analysis of raw accelerometry data and provide insights into potential solutions. In particular, we describe the size and complexity of the data, the within- and between-subject variability and the effects of sensor location on the body. We also provide a short tutorial for dealing with sampling frequency, device calibration, data labeling and multiple PA monitors synchronization. We illustrate these points using the Developmental Epidemiological Cohort Study (DECOS), which collected raw accelerometry data on individuals both in a controlled and the free-living environment.

## 1 Introduction

Wearable physical activity (PA) monitors provide detailed, continuous, and objective measurements of individual PA in the free living environment. They can complement or completely replace current subjective measurements collected via questionnaires. Recent advances in technology and the decreasing cost of wearable devices led to an explosion in the popularity of wearable technology in health research. Here we argue that, just like any new measurement used in health science, there is a need to understand, reproduce, and communicate the measurements produced by these new devices. This can lead to improved design of experiments, higher quality of the acquired data, and more generalizable results. At the core of all modern PA monitors there is a small accelerometer, a Microelectromechanical system (MEMS) that measures accelerations relative to the Earth’s gravitational field. Hence, PA monitors are often referred to as wearable accelerometers. The output of these devices is a three-dimensional time series of accelerations expressed in gravitational units in the frame of reference of the device. More clearly, the device has its own frame of reference up-down, left-right, backward-forward. This frame is typically different from and can change with the frame of reference of the person who wears the device or who observes the experiment. These raw data produced by accelerometers are transformed using various algorithms into PA summaries, which have different labels (e.g. steps, calories, activity counts) and can be aggregated at different temporal resolutions (e.g. minutes, hours, or days).

Due to battery and memory limitations, PA monitors used to return only aggregated minute-level data in the form of proprietary activity counts (AC) (Chen et al, 2012). The definition of ACs varies between- and within-device manufacturers, across time, body location and between studies. Despite these initial problems, they have been used effectively as a relative measure of PA within the same study, especially when the same devices and software were used and devices were calibrated (Matthews et al, 2008; Healy et al, 2011; Schrack et al, 2014; Xiao et al, 2015). As battery and memory limitations have been mitigated, it has become possible to collect and store high-throughput, three-axial, sub-second level acceleration data, ranging between 10 and 200 observations per second. PA monitors have also become increasingly sophisticated and are now routinely equipped with a selection of supplementary sensors including gyroscopes, thermometers, inclinometers, pulsometers, light intensity and skin conductance sensors. Data collected by these supplementary sensors are beyond the scope of this paper.

The collection of raw accelerometry data opens a spectrum of new scientific and analytic problems. For example, researchers do not need to rely on proprietary aggregated measures and can use well-defined, open-source, reproducible summaries of the data. This allows to compare and combine studies that collect raw accelerometry data at the same location on the body and provides explicit measures of activity on a recognized measurement scale. For example, UKBiobank (Doherty et al, 2017) uses the magnitude of the 3-dimensional vector of acceleration summarized in 5-second intervals, whereas the Women’s Health Initiative (WHI) study has explored 1-second summaries based on standard deviations of accelerometry along each axis (Bai et al, 2016). Raw and summarized data have also been used for recognition of activity types. A common approach is to derive accelerometry data features in a particular window and use supervised classification approaches to predict activity type (Pober et al, 2006; Staudenmayer et al, 2009; Attal et al, 2015). Extensive reviews of classification techniques for activity recognition from accelerometry data are provided by Bao and Intille (2004) and Preece et al (2009). Dictionary learning based on raw accelerometry Bai et al (2012); He et al (2014); Xiao et al (2016) has also been proposed in the context of fine-resolution movement prediction. The increased granularity of the sub-second level data may contain important additional information, but it also creates new challenges. Indeed, the volume and structure of the data are much more complex for the raw data, especially when it is recorded for weeks at a time in free-living settings. In this paper we discuss problems related to the collection and analysis of sub-second level accelerometry data. To illustrate these problems, we use data collected as a part of Developmental Epidemiological Cohort Study (DECOS) in a controlled and in the free-living environment.

Some of these problems have been well documented in the literature. For example, Trost et al (2005) highlighted problems associated with device se-lection, placement of accelerometers, epoch length, and compliance enhancing strategies. Schrack et al (2016) discussed the limitations of using uncalibrated, population-level data extraction algorithms in older adults. Staudenmayer et al (2012) and Troiano et al (2014) noted that the diversity of measurements and analytic methods for accelerometer data makes it difficult to compare results across different studies, and advocate for standardization of measurement and pre-processing pipelines across studies.

Currently, there is no universally accepted and standardized approach for measuring PA using wearable accelerometers in health research. However, some excellent guidelines and standardized protocols have been published. For example, Matthews et al (2012) compiled a list of best practices for making decisions about important choices, such as the number of monitors needed, device placement, device initialization, device tracking, and data collection. They also provide guidance on how to report the use of PA monitors in populationbased studies. Freedson et al (2012) provided recommendations for the use of wearable monitors for assessing PA for researchers, end users, as well as developers of activity monitors. The authors also provide guidelines for sensor output calibration and validation and discuss necessary steps for maximizing the generalizability of the data analysis.

Unfortunately, harmonization of PA data between existing studies is often impossible due to the different formulation and interpretation of activity counts, body placement, and lack of universally accepted measurement. For example, step counts produced by Fitbit cannot be compared to activity counts produced by ActiGraph, as they are not even on the same scale. In fact, is has been shown that step counts might differ substantially when measured by different devices (Storti et al, 2008; Fortune et al, 2014). Activity counts measured by the same device may also be different depending on the sensor location (Fairclough et al, 2016). For example, a device located on the hip or ankle is sensitive to ambulation, whereas a device located on the wrist will detect both ambulation and hand movements. Moreover, some activity counts may change substantially with the sampling frequency (Brønd and Arvidsson, 2016) even for the same device placed at a particular body location. These problems are not likely to be resolved as long as summary data obtained via proprietary manufacturer algorithms continue to be used. The best strategy is to go back to the raw data and construct open-source data pre-processing approaches that become increasingly accepted by the community. This, unfortunately, is not a panacea, as pre-processing pipelines need to take into account problems associated with device calibration, missing data, data size and complexity, and measurement translation and communication.

To provide a concrete illustration of these problems and some potential solutions, we use the Developmental Epidemiological Cohort Study (DECOS) data, which is described in Sections 2.1-2.2. Next we introduce the notation and statistical methods used to pre-process and summarize the data in Section 2.3. In Section 3 we illustrate issues related to the analysis and interpretation of raw accelerometry data and review some published approaches designed to handle the complexity and heterogeneity of accelerometry data collected in the free-living environment. In Section 4 we summarize the ideas and discuss their implication for health studies using raw accelerometry data.

## 2 Methods

### 2.1 Study participants

Forty-nine community-dwelling older adults were recruited from the Pittsburgh, Pennsylvania area for the Developmental Epidemiologic Cohort Study (DECOS), part of the National Institute on Aging (NIA) Aging Research Evaluating Accelerometry (AREA) project (Lange-Maia et al, 2015). DECOS is a cross-sectional study designed to examine the impact of accelerometry wear location on PA and sedentary behavior assessment among healthy older adults. Individuals were excluded from DECOS if they suffered from any of the following conditions: hip fracture, stroke in the past 12 months, cerebral hemorrhage in the past 6 months, heart attack, angioplasty, heart surgery in the past 3 months, chest pain during walking in the past 30 days, current treatment for shortness of breath or a lung condition, usual aching, stiffness, or pain in their lower limbs and joints and bilateral difficulty bending or straightening the knees fully.

### 2.2 Data collection

Participants were equipped with three tri-axial wearable PA monitors (Acti-Graph GT3x+) that collected raw accelerometry data at a sampling frequency of 80Hz (80 observations per second for each axis). Monitors were located on the hip using an elastic belt and on both wrists using watch straps. During the “in-the-lab” phase of the experiment all participants were asked to perform a series of physical tasks including: lying still, standing still, washing dishes, sitting still, dough kneading, dressing, folding towels, vacuuming, shopping, writing, dealing cards, standing up from a chair, walking for 20 meters, walking for 20 meters with arms crossed on the chest, fast walking for 20 meters, fast walking for 20 meters with arms crossed on the chest, treadmill walking at 1.5mph for 5 minutes, walking for 40 meters and fast walking for 400 meters. Before each task participants were given verbal instructions by a supervising trained professional recording the times of the beginning and end of each task with a stopwatch. During the free-living portion of the experiment, participants were equipped with accelerometers for seven consecutive days and were told to maintain their normal, unsupervised, free-living activities. They were instructed to take off the activity monitors only during sleep.

### 2.3 Open-source summaries of accelerometry data

The raw accelerometry data are collected along three orthogonal axes in the device-specific frame of reference. We denote the vector of raw acceleration data by {**x**(*t*) = *x*_1_(*t*), *x*_2_(*t*), *x*_3_(*t*), where *x*_*m*_(*t*)} is the acceleration measurement along the *m* (= 1, 2, 3) axis at time *t* (for notational simplicity we drop the subject index). The acceleration time series, **x**(*t*), are sampled at a fixed frequency *f*. For example, in our application, the sampling frequency is 80 Hz, which means that for each second along each axis there are 80 acceleration recordings. Here we describe four open-source methods that have been used in the literature to provide summaries of accelerometry data. All methods consider non-overlapping time windows of a given length *H* and reduce the 3*H* measurements in the window to a single number.

Euclidean Norm Minus One (ENMO) was first introduced as a summary metric for accelerometry data in van Hees et al (2013). It is directly based on the Euclidean norm of **x**(*t*), defined as

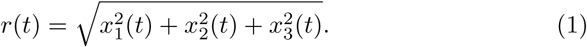

The ENMO at time *t* is defined as *r*(*t*) −1 when *r*(*t*) −1 ≥ 0 and 0 otherwise, or notationally, max [*r*(*t*) −1, 0]. Further, the ENMO in a window of size *H* is defined as the average ENMO across the time points in that window. Formally,

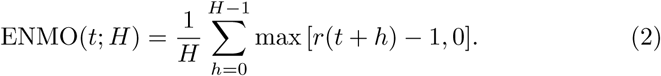

ENMO can be quite sensitive to calibration errors when the device-specific ENMO at rest is not close to zero. An additional calibration procedure was introduced to mitigate the effects of calibration (van Hees et al (2014)). This procedure performs a linear transformation on the raw data before computing the Euclidean norm, resulting in a new version of ENMO. Calibration para-meters *a*_*m*_ and *d*_*m*_ are estimated for each axis *m* = 1, 2, 3, which are later used to linearly transformed the original data to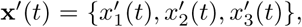, such at 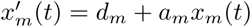 for *m* = 1, 2, 3.

The Vector Magnitude Count (VMC) is an aggregation statistic that is also known as the Mean Amplitude Deviation (MAD) Vähä-Ypyä et al (2015). We use the notation VMC to avoid the confusion between the two MAD acronyms used in accelerometry literature, one for mean amplitude deviation (Vähä-Ypyä et al, 2015) and one for median amplitude deviation (Mariani et al, 2013). VMC computes the ℒ_1_ norm in each time window *H*. Denote the average Euclidean norm in the window of length *H* starting at *t* as 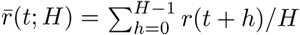 Then VMC is defined as

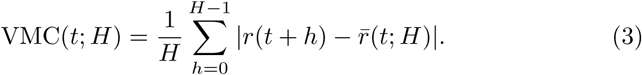

The unnormalized Activity Index (AI_0_) (Bai et al (2014) is a measure ba-sed on the combination of the three within-axis standard deviations of the raw accelerometry signal. Because AI_0_ subtracts the local mean of the accelerometry signal, calibration is intrinsic and local, which allows it to adapt to cases when the device is not calibrated, when it gets decalibrated during studies, or when the device exhibits time-dependent decalibration. Let *σ*_*m*_(*t*; *H*) be the standard deviation of the acceleration along axis *m* = 1, 2, 3 in the window of length *H* starting at *t*. The exact formula is

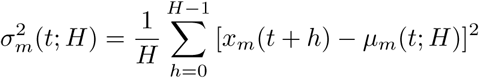

where 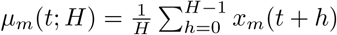. Then formally,

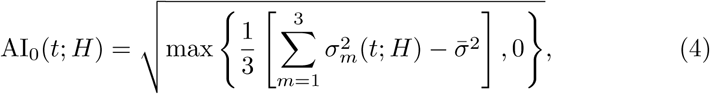

where 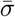 is the systematic noise standard deviation calculated using the data collected during some non-moving period. The unnormalized Activity Index (AI_0_) is expressed in Earth gravitational units.

The normalized Activity Index (AI, Bai et al (2016)) is strongly related to the unnormalized Activity Index (AI_0_). The only difference is that the axis-specific variances are divided by the device-specific systematic noise. More specifically, AI is defined as follows

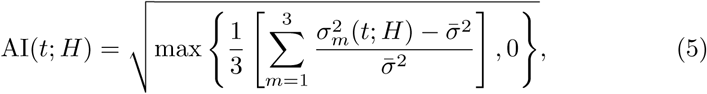

The downside of using AI versus AI_0_ is that its scale is no longer in Earth gravitational units. Instead it is expressed in multiples of noise standard deviation. The advantage of AI could be when the devices are not calibrated in terms of their noise level at rest, which may induce batch effects. This, however, seems to be a smaller problem than the internal calibration obtained by subtracting the local mean in AI and AI_0_.

In the remainder of this paper, we use ENMO, VMC and AI_0_. We do not use normalized Activity Index (AI) so as to keep presentations of the statistics measurements on the same scale.

## 3 Statistical challenges and examples

### 3.1 Data volume and complexity

Figure 1 displays three time series that represent the acceleration along each of the three orthogonal axes of an accelerometer located at the wrist. The top panel of Figure 1 presents 24 hours of data, with each axis data shown in a different color. In this example, the first observation was taken at 12AM, while the last was taken 24 hours later, also at 12AM. The middle panel in Figure 1 displays a particular one hour interval from 8 : 40AM to 9 : 40AM (indicated in the top panel as a dashed-line rectangle). The bottom panel displays the one minute interval marked as a dashed-line rectangle in the middle panel (from 8 : 51AM to 8 : 52AM). The signal was acquired at a sampling frequency of *f*_*s*_ = 80Hz. Therefore, the number of observations per subject quickly explodes. For example, a week of accelerometry data collected at 80*Hz* results in 80 * 60 * 60 * 24 * 7 = 48, 384, 000 observations for each of the three axes. Thus, even for a small multi-subject observational study researchers are faced with datasets consisting of billions of observations. This enormous volume of data creates challenges at every level of the scientific investigation. Storage and operational memory of modern computers is not unlimited and well-optimized solutions are needed for data management. Conducting exploratory data analysis, visualization, and modeling requires additional computational and methodological resources. Therefore, it is essential to implement carefully planned protocols for collection, management and analysis of the data. In recent years several protocols and experimental design guidelines have been proposed (Esliger et al, 2005; Cain, 2014; NHANES, 2011), though an universally accepted approach is still elusive. This could be due to the constant change of the technological and methodological landscape. For example, the US NHANES survey protocols have changed between survey cycles 2003-2004 (NHANES, 2006), 2005-2006 (NHANES, 2008), 2011-2012 and 2013-2014 (NHANES, 2011) both in terms of device type and body location. In the 2003-2004 and 2005-2006 cycles, participants wore an ActiGraph 7164 on a waist belt for 7 days during the non-sleeping time. In later cycles, the GT3X+ ActiGraph waterproof model was used on the non-dominant wrist for 7 days without taking it off. The protocol change was reported (Troiano et al, 2014) and was designed to improve participant compliance.

**Figure 1.**
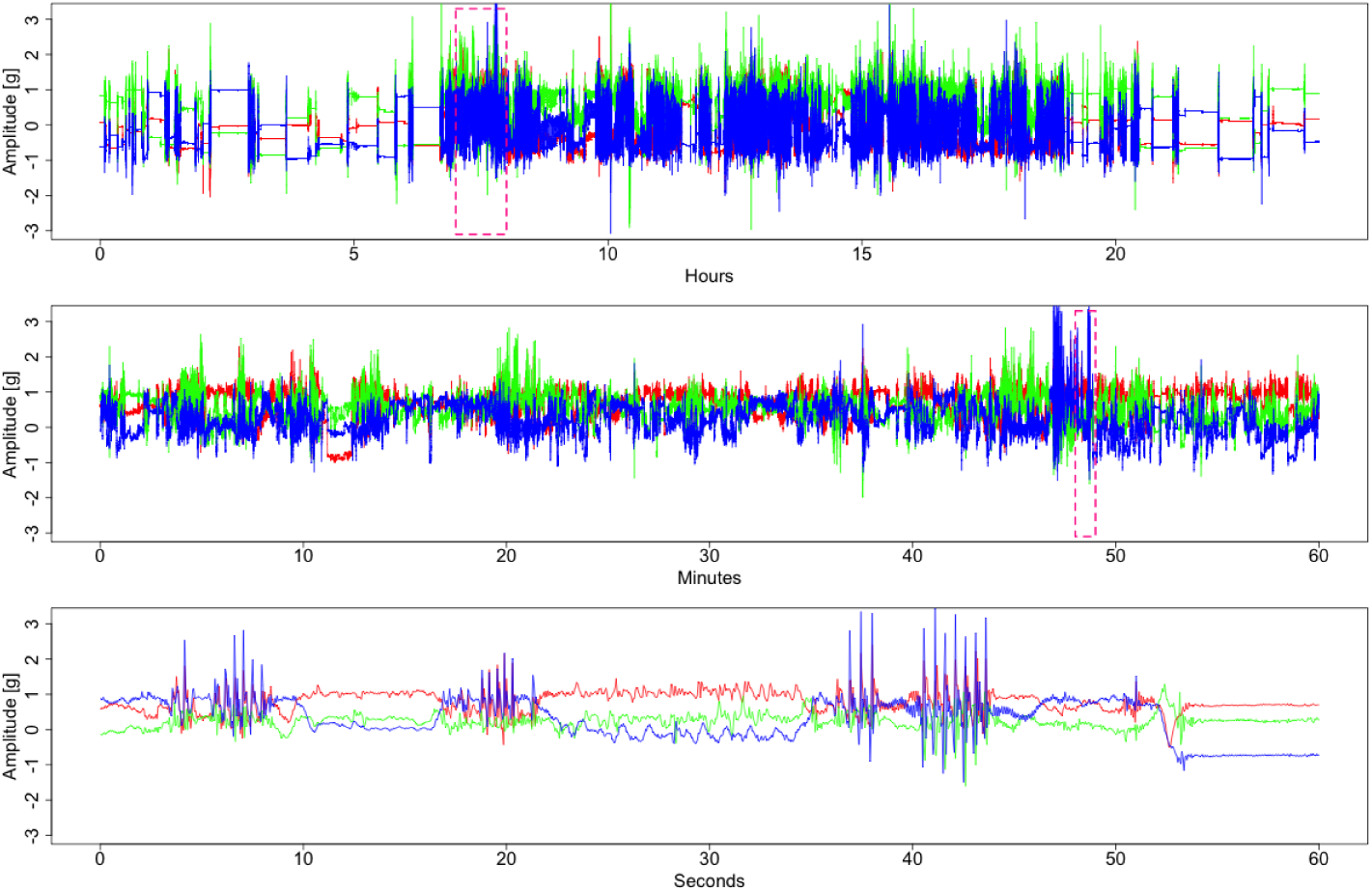
Acceleration values from three orthogonal axes of an accelerometer located on the left wrist. Each axis data is shown in a different color. The top panel displays 24 hours of data collected between 12AM and 12AM. The middle panel displays a one hour interval from 8 40AM to 9 40AM (indicated in the top panel as a dashed-line rectangle). The bottom panel displays a one minute interval from 8 51AM to 8 52AM marked as a dashed-line rectangle in the middle panel. The signal was acquired at a sampling frequency of *f_s_* = 80Hz.

The current approach to reduce size and complexity of the raw accelerome-ter data is to create aggregated summaries in fixed time intervals, as described in Section 2.3. While these summaries reduce the volume of data, the potential loss of information can be substantial. To recover some of the information, a few methods have been proposed for recognition of activities using raw accelerometry data. Some approaches focus on prediction of a group of movement types (Lyden et al, 2014), whereas others focus on prediction of a specific movement type (e.g. walking) (Urbanek et al, 2015). Characterizing the kinematics of human walking both in the lab and in the free-living environments using accelerometry data could provide previously unavailable information on the physical condition of individuals (Studenski et al, 2011).

### 3.2 Data heterogeneity

Interpretation of raw accelerometry data is an open and challenging problem due to the high heterogeneity of data, both within- and between-subjects. Within-subject variability is observed when one person performs the activity, but the characteristics of that activity change. For example, when walking, consecutive strides differ slightly in duration and shape due to natural stride-to-stride variability (IJmker and Lamoth, 2012; Urbanek et al, 2017). They can also differ substantially during the day, depending on the level fatigue of the individual, context of walking (e.g. hiking versus shopping), and local constraints (e.g. running to a meeting when late versus slow walking to the kitchen in the morning). Between-subject variability contains additional factors due to differences in body size, musculature, will, and ability to perform certain tasks. To further illustrate these points, the left column of Figure 2 displays acceleration data collected on the left wrist during walking for two individuals. The periodic character of time series is striking. However, both the duration and amplitude of accelerometry data can vary from cycle to cycle. A more extreme example is displayed in the right column of Figure 2, where the same two individuals perform the getting dressed activity. Indeed, the more random character of the observations and lack of synchronization between- and within individuals is remarkable.

**Figure 2.**
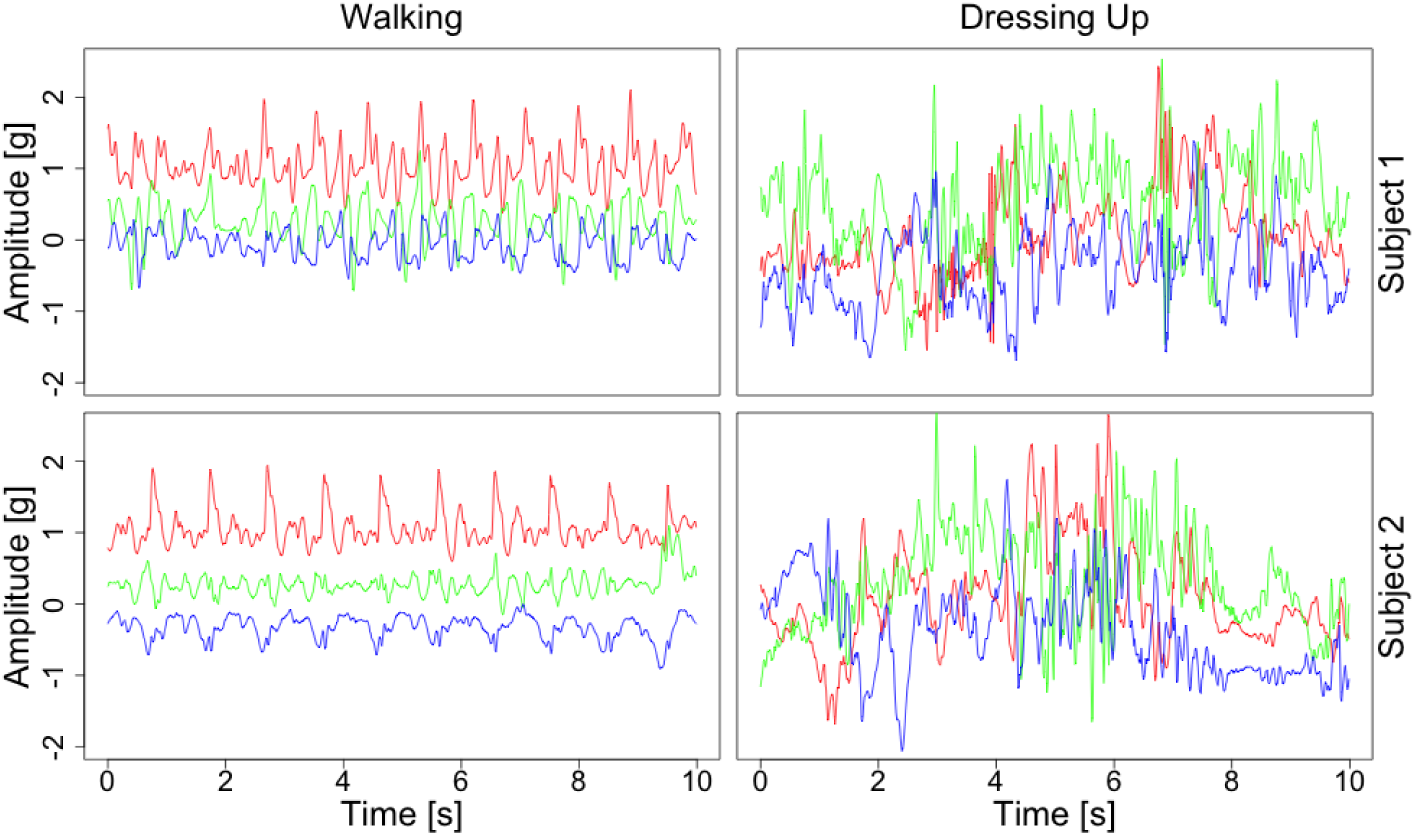
Data recorded by an accelerometer located on the left wrist while walking (left column) and getting dressed (right column), for two individuals (top and bottom row). Each axis is shown in a different color

These data indicate that it is important to better define the types of physical activity. Indeed, the bottom panels of Figure 2 show that data can be very different even for what is defined as the same type of activity (e.g. getting dressed). In retrospect, this should not be surprising as “getting dressed” is a complex task that is only vaguely defined, may involve different types of clothes, body sizes and shapes, movements that individuals use to getting dressed, and order of various tasks. It should be apparent that if we want to make any progress in this area, we need to identify well-defined sub-movements that then translate them into research language. Clearly, in this case, the activity “getting dressed” does not have a sharp definition, especially from the point of view of an accelerometer. Things may be different if we had a video camera instead of an accelerometer, but this raises other problems that exceed the scope of the current paper.

Classifying activity types while accounting for between- and within-subject variability is under intense methodological development. The approach discussed in Bai et al (2012) and He et al (2014) uses short segments of training accelerometry data, called “movelets”, to construct dictionaries used as reference to predict activity types on new data. Dictionaries are activity- and subject-specific to account for the individual variations in movement patterns across subjects. Xiao et al (2016) proposed a related movelet-based method to use labeled activity data from some subjects to predict the activity labels of other subjects.

### 3.3 Sensor location

The location of the accelerometer sensor on the body can also have substantial effects on predicting activity types, PA volume, and PA distribution as a function of time of the day. Indeed, the sensor collects only its own acceleration, which is a proxy of the acceleration of the particular body location where the sensor is attached. Therefore, a wrist sensor will produce different signals from a hip or thigh sensor. To illustrate this point, Figure 3 displays the raw accelerometry data collected simultaneously by sensors located on the hip (left column) and left wrist (right column) while dealing cards (top panel), getting dressed (middle panel) and walking (bottom panel). For dealing cards, an activity that requires mostly hand-movements, the signal amplitude for the wrist sensor is much higher than for the hip sensor. For more complex, whole-body activities, such as getting dressed, the signals corresponding to both locations have higher amplitudes than for dealing cards. However, the amplitude of the signal at the wrist is higher and there are no clear correlations between the hip and wrist signals. In the case of walking, the amplitudes of the data collected on the wrist and the hip look periodic and highly correlated. This is likely due to the fact that both hands and legs are involved in walking, with roughly the same frequency of movement. The slightly lower amplitudes observed at the wrist are probably due to the more intense PA in the lower body during walking. In our experience, an accelerometer placed at the ankle would display even higher amplitudes of the acceleration signal.

**Figure 3.**
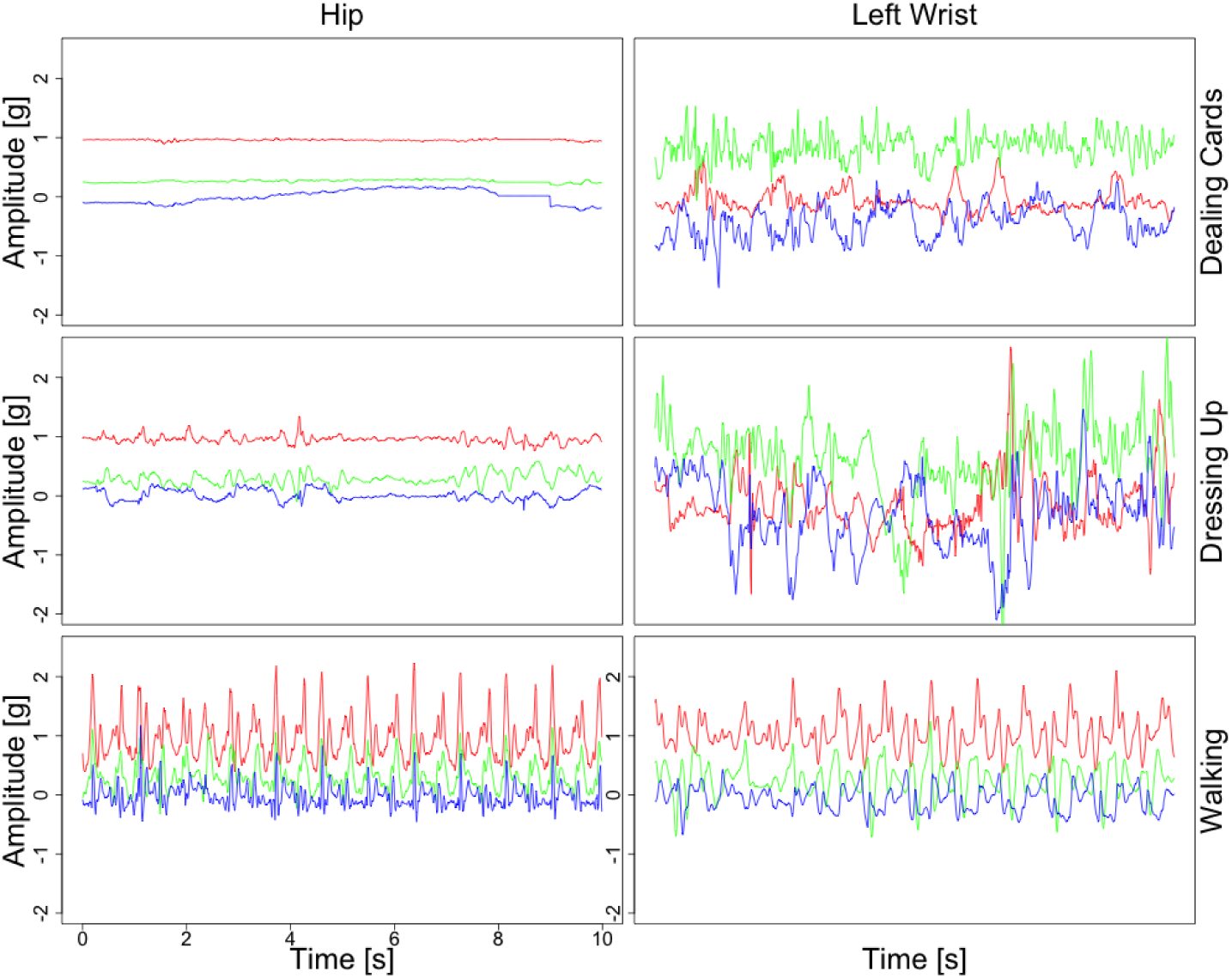
Acceleration from three orthogonal axes of an accelerometer located on the hip (left column) and left wrist (right column), while dealing cards (top row), getting dressed (middle row) and walking (bottom row). Each axis data are shown in a different color.

Figure 4 provides the summary metrics introduced in Section 2.3 using data collected during various activities performed in the controlled lab environment. The boxplots for ENMO, VMC and AI_0_ are displayed in the top, middle and bottom panels, respectively, while the data for the hip and left wrist are displayed in the left and right columns, respectively. All summary statistics are calculated for a window size of 5 seconds during writing, washing dishes, vacuuming, getting dressed and walking for each of 5-second intervals and all 49 individuals. The difference between the wrist and hip data can be observed for all summaries and activities requiring dynamic upper body movements (washing dishes, vacuuming). These differences are much lower for low intensity hand movements (writing), though AI_0_ seems to better capture these differences. Also, substantial differences can be observed for moderate-intensity whole-body activities (getting dressed), where, again, AI_0_ seems to be more sensitive. For walking, the difference in the distribution of all summaries between the wrist and the hip is relatively small. When comparing AI_0_ versus ENMO versus VMC at the wrist there is a more clear differentiation between activities. This is likely due to the fact that AI_0_ automatically corrects for possible device miss-calibration, whereas the other methods do not. It is an open problem whether the methods perform more similarly after device and/or signal calibration.

**Figure 4.**
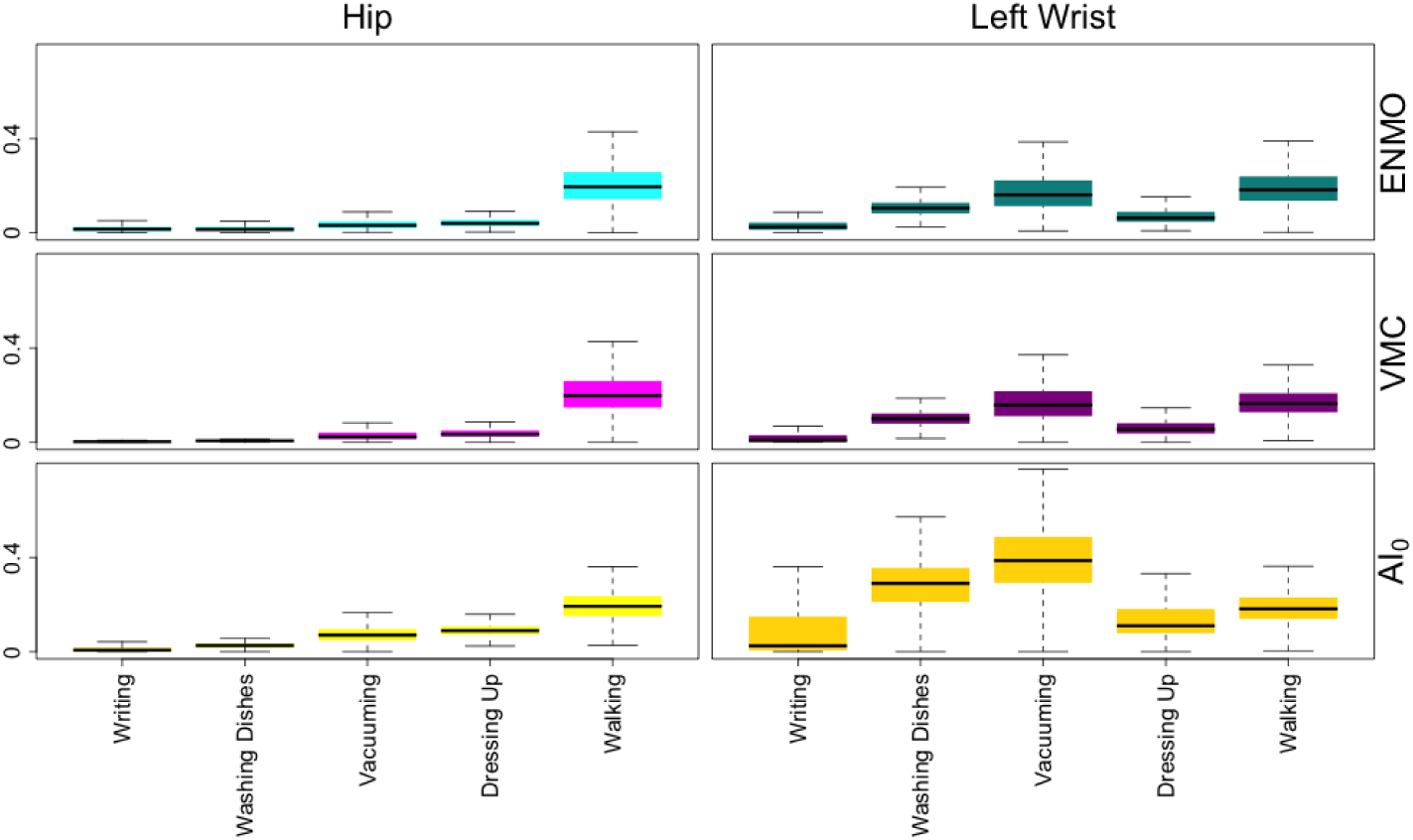
Boxplots of ENMO, (top panels) VMC (middle panels) and AI_0_ (bottom panels) statistics derived for *τ* = 5 seconds-length intervals of data collected from the hip (left column) and left wrist (right column) during writing, washing dishes, vacuuming, getting dressed and walking (x-axis), for all 49 individuals.

**Figure 5.**
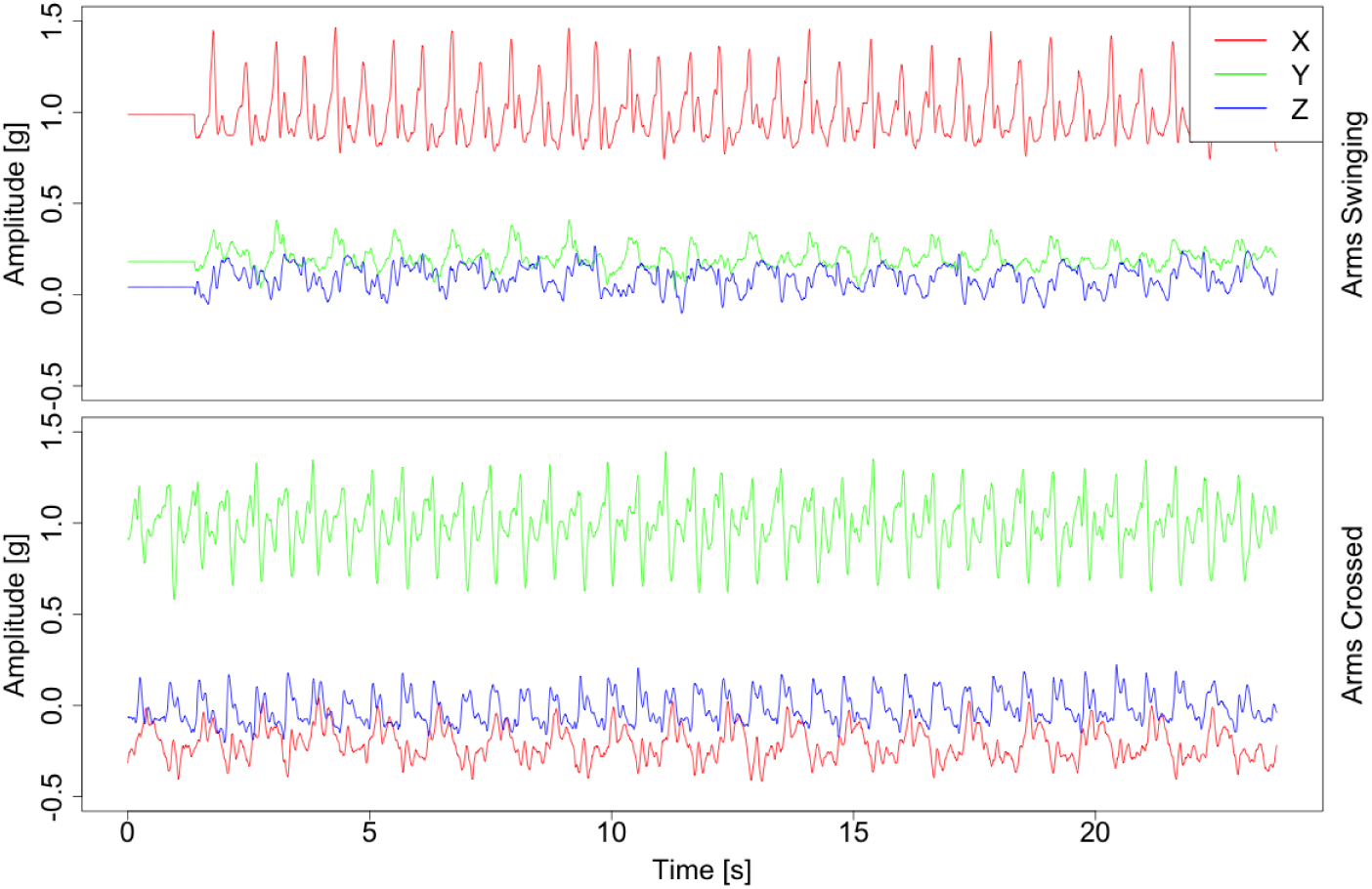
Acceleration values from three orthogonal axes of an accelerometer located on the left wrist, collected during two walking tasks performed by the same individual. Each axis data is shown in a different color. The upper panel corresponds to walking with both hands moving naturally, whereas the bottom panel corresponds to walking with arms crossed on the chest.

Summaries have been used extensively in the literature. For example, Koster et al (2016) showed that different cut-points of the vector magnitude can be used for the classification of sedentary time in older adults using both hip-worn and wrist-worn ActiGraph accelerometers. It has also been shown that activity recognition algorithms perform differently across body locations. For example, Rosenberger et al (2013) reported greater sensitivity and specificity of both sedentary and moderate- to vigorous-intensity PA when using accelerometry data collected at the hip compared to wrist. Trost et al (2014) reported higher accuracy for hip-derived than for wrist-derived data in the classification of specific, whole-body engaging activities. Higher accuracy of classifying sitting was obtained with data collected from the wrist compared to data collected from the hip. Del Din et al (2016) showed that estimated gait characteristics, such as step time and length, can depend on body location and suggested that the chest location is more appropriate than the wrist. We conclude that the body location may lead to different results, that body location will favor movements that directly engage that particular location, and that translating various summaries into well defined activity categories (e.g. walking, sedentary) requires location-specific calibration of the accelerometry summaries.

### 3.4 Device rotation

Because wearable PA monitors collect acceleration data relative to earth gravity, they are sensitive not only to movement but also to their own orientation with respect to Earth’s gravity. To better understand this, the top panels in Figure 5 display accelerometry data collected by a wrist-worn device during two walking tasks performed by the same individual. The upper panel corresponds to walking with both hands moving naturally, whereas the bottom panel corresponds to walking with arms crossed on the chest. The change in the device orientation is manifested in the change of mean values, most clearly seen in the signals shown in green. In the free living environment changes in device rotation are quite common and can be due to multiple sources. For example, when walking individuals could sway their hands normally, hold them in their pockets or perform an activity (e.g. holding a smart phone), walking with hands hanging loose or with hands in the pockets. Additionally, the device can rotate around the wrist or move higher or lower on the hand, resulting in an altered distribution of the observed signal.

To prevent a device from rotating, it can be directly attached to the skin with adhesive pads or affixed indirectly with a waistband clip or elastic belt (Matthews et al, 2012). Indeed, it has been recommended that devices should be fitted as tightly to the body as possible (Boerema et al, 2014). However, even with these precautions, belts can loosen up, resulting in device orientation changes (O’Neill et al, 2017). Devices that adhere to the skin might be attached upside-down or placed in a slightly different position when detached and reattached. Moreover, for devices that have it, the orientation chart can be obscured, which can result in improper device orientation (Edwardson et al, 2016). We consider that all these precautions should be carefully implemented and adapted to the specific problem that one tries to address. In addition, it is important to understand the extent of the problem in specific applications and, when possible, attempt to correct it. Several methods were proposed to address this problem by rotating the observed three-dimensional vector to the common, reference, orientation (Xiao et al, 2016; Yurtman and Barshan, 2017). When one uses summary metrics that are robust to device orientation, this problem is less important. For example, ENMO, VMC and AI are all rotation invariant.

### 3.5 Sampling frequency

Sampling frequency *f*_*s*_ (expressed in Hz) is the parameter describing how often accelerometry data are collected by the device. In modern wearable accelerometers sampling frequency usually ranges between 10 to 200 Hz, though it could be set as high as 1000 Hz for specialized applications, such as precise human movement tracking during sport activities (Dominguez-Vega et al, 2015). One of the hidden problems is that the summaries produced by various devices can depend on the sampling frequency of the device. This could have substantial implications if, for example, in a study the sampling frequency is varied between- and within-individuals and/or devices. For example, according to ActiGraph, LLC, the manufacturer of ActiGraph PA monitors, the observed activity counts depend on the sampling frequency (ActiGraph, 2016).

Figure 6 illustrates the boxplots of ENMO (top panel), VMC (middle panel) and AI_0_ (bottom panel) calculated for all 49 participants during writing, washing dishes, vacuuming, getting dressed and walking in 5-second time intervals. Data were collected by the device located on the left wrist with the original sampling frequency *f*_*s*_ = 80 Hz. Additionally, data has been decimated to simulate sampling frequencies of 40, 20 and 10 Hz. The median summary values are reported in Table 1. Interestingly, all open source summaries are relatively stable as a function of frequency, with stronger decreases at 10 Hz. For the AI_0_ statistic, the change is most pronounced for walking, where the median AI_0_ decreases by 19.0% from 80 to 10 Hz. For the other activities the reduction is much smaller in the 1 to 7% range. These smaller differences in open source measures are encouraging. Indeed, Brønd and Arvidsson (2016) investigated the effects of sampling frequency on ActiGraph activity counts. They compared results obtained during walking and running with the default 30 Hz sampling frequency with 40 and 100 Hz sampling frequencies. For fastrun activity, they reported approx. 6,800 counts per minute (cpm) mean for default 30 Hz frequency, and an increase of mean cpm as high as 24% for 40 Hz and 18% for 100 Hz frequency.

**Figure 6.**
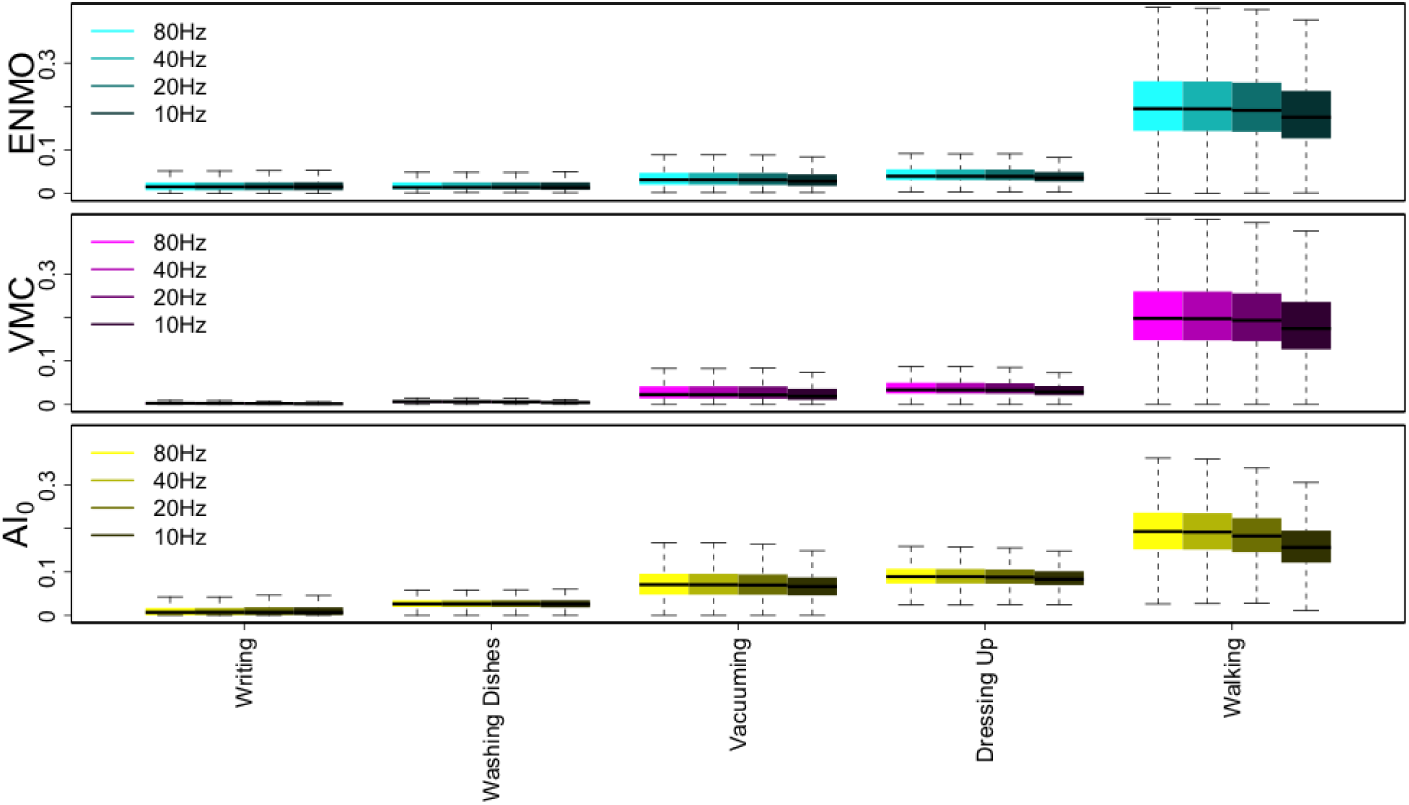
Boxplots of ENMO, (top panels) VMC (middle panels) and AI_0_ (bottom panels) for 5 second time windows. Data is shown for all 49 individuals in the study and were collected from the left wrist during writing, washing dishes, vacuuming, getting dressed and walking (x-axis). Data were collected with the original sampling frequency *f_s_* = 80 Hz and then decimated to simulate sampling frequencies of 40, 20 and 10 Hz.

**Table 1.** Median values of ENMO, VMC and AI_0_ statistics derived for *τ* = 5 seconds-length intervals of data collected from the left wrist during writing, washing dishes, vacuuming, getting dressed and walking, for all 49 individuals. Results are obtained for original sampling frequency *f_s_* = 80*Hz* (3rd column) and simulated sampling frequencies *f_s_* = 40, 20, 10 *Hz* (4-6th columns).

| Statistic | Activity | 80Hz | 40Hz | 20Hz | 10Hz |
| --- | --- | --- | --- | --- | --- |
| ENMO | Writing | 0.015 | 0.015 | 0.016 | 0.016 |
| ENMO | Washing Dishes | 0.014 | 0.014 | 0.014 | 0.014 |
| ENMO | Vacuuming | 0.031 | 0.031 | 0.031 | 0.028 |
| ENMO | Getting Dressed | 0.040 | 0.040 | 0.040 | 0.036 |
| ENMO | Walking | 0.195 | 0.195 | 0.191 | 0.176 |
| VMC | Writing | 0.003 | 0.002 | 0.002 | 0.002 |
| VMC | Washing Dishes | 0.007 | 0.006 | 0.006 | 0.004 |
| VMC | Vacuuming | 0.023 | 0.022 | 0.022 | 0.019 |
| VMC | Getting Dressed | 0.034 | 0.033 | 0.033 | 0.029 |
| VMC | Walking | 0.198 | 0.197 | 0.193 | 0.175 |
| $AI_0$ | Writing | 0.007 | 0.007 | 0.007 | 0.007 |
| $AI_0$ | Washing Dishes | 0.026 | 0.027 | 0.027 | 0.026 |
| $AI_0$ | Vacuuming | 0.071 | 0.071 | 0.070 | 0.066 |
| $AI_0$ | Getting Dressed | 0.089 | 0.089 | 0.088 | 0.083 |
| $AI_0$ | Walking | 0.193 | 0.192 | 0.183 | 0.156 |

The Table 3 in Appendix A) provides the medians, 25-th, and 75-th percentile for the same five activities, but broken down by subject for 5 different subjects. Results are only shown for the hip accelerometer, though similar results are available for wrist accelerometers. The ranking of activity intensities by medians across subjects is the same, but the medians for individual subjects are quite variable even for the same activity and summary metric. For reference, at the hip, AI_0_ is roughly around 0 milli g for writing, between 1 and 2 milli g for washing dishes, 3 and 14 milli g for vacuuming, 5 and 10 milli g for getting dressed, and 20 to 100 milli g for walking. Here we used only the minimum of the first and maximum of the third quartiles across the five subjects to create these ranges. These measures are averages per second during the 5 second intervals and not totals. If one would like to transform these numbers into totals per minute then the values need to be multiplied by 60; similar for other intervals.

### 3.6 Measurement bias and batch effects

Measurement bias is the difference between the measured accelerations and their true values. Estimation of both bias and measurement error of accelerometry data requires a dedicated experimental setup utilizing calibrated vibration exciters. The device is exposed to a known acceleration and the measured acceleration time series is compared to this known acceleration, as described in Bassett et al (2012). Authors noted that newer devices, such ActiGraph GT1M, undergo initial unit calibration during production and are supposed to be calibrated for as long as they are used. However, the calibre tion standards of the different manufacturers may be different. Therefore, we recommend doing some basic calibration checks before utilizing the device in a study.

Even when dynamic calibration is conducted, bias in static measurements can still exist. Ideally, an accelerometer resting on a flat surface, with one measurement axis oriented perpendicularly to the ground, should measure constant acceleration of 1*g* for that axis and 0*g* for the other two orthogonal axes. In practice, measurements may deviate slightly due to their imprecision or to the quality of assembly, which may, for example, misalign the accelerometer axes with the monitors’ casing. In theory, the vector magnitude (Eq. (1)) for resting state (no movement) is equal to 1*g*, as the earth’s gravity is the only force acting on the accelerometer. In practice, however, the vector magnitude at rest can be slightly different from 1*g*. To quantify this calibration bias, we calculated a vector magnitude *r*(*t*) averages from 5-second time windows, denoted 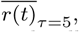, for the acceleration signal collected from the hip during sitting still activity, for all 49 subjects. We report the percentiles of this distribution in Table 2. Ideally, if no bias was present in the data, 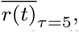 values should all be equal to 1. The median is 1.007 indicating very close agreement with what we expect. However, the 5 and 95 percentiles were 0.964 and 1.047, indicating that 10% of the devices have a deviation of 5% or more from 1g at rest.

**Table 2.** Percentiles of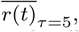, vector magnitude *r*(*t*) averages from 5-second time windows, for the acceleration signal collected from the hip during sitting still activity, for all 49 subjects.

| Percentile | 5 | 25 | 50 | 75 | 95 |
| --- | --- | --- | --- | --- | --- |
| $\overline{r(t)}_{\tau=5} [g]$ | 0.964 | 0.991 | 1.007 | 1.016 | 1.047 |

**Table 3.** Summary of the four statistics: ENMO, VMC, AI_0_ and AI for five selected subjects and all subjects: median, 25-th percentile and 75-th percentile (percentiles are reported in brackets), obtained from accelerometry data collected at the hip during five activities: writing, washing dishes, vacuuming, getting dressed and walking.

| id | Writing | Washing Dishes | Vacuuming | Getting Dressed | Walking |
| --- | --- | --- | --- | --- | --- |
| **ENMO** |  |  |  |  |  |
| Sub. 1 | 0.021<br>(0.019,0.023) | 0.033<br>(0.032,0.033) | 0.037<br>(0.035,0.039) | 0.035<br>(0.032,0.038) | 0.189<br>(0.161,0.204) |
| Sub. 2 | 0.084<br>(0.083,0.085) | 0.029<br>(0.028,0.030) | 0.031<br>(0.027,0.047) | 0.055<br>(0.052,0.058) | 0.147<br>(0.122,0.158) |
| Sub. 3 | 0.024<br>(0.022,0.024) | 0.037<br>(0.037,0.038) | 0.039<br>(0.037,0.043) | 0.064<br>(0.059,0.071) | 0.175<br>(0.157,0.198) |
| Sub. 4 | 0.005<br>(0.003,0.006) | 0.008<br>(0.006,0.009) | 0.043<br>(0.028,0.089) | 0.041<br>(0.024,0.051) | 0.181<br>(0.154,0.192) |
| Sub. 5 | 0.017<br>(0.014,0.023) | 0.019<br>(0.018,0.019) | 0.038<br>(0.030,0.066) | 0.035<br>(0.031,0.044) | 0.297<br>(0.263,0.318) |
| **All Sub.** | **0.015**<br>**(0.007,0.025)** | **0.014**<br>**(0.009,0.025)** | **0.031**<br>**(0.020,0.047)** | **0.040**<br>**(0.031,0.055)** | **0.195**<br>**(0.143,0.257)** |
| **VMC** |  |  |  |  |  |
| Sub. 1 | 0.004<br>(0.002,0.005) | 0.006<br>(0.006,0.007) | 0.020<br>(0.015,0.023) | 0.021<br>(0.017,0.025) | 0.194<br>(0.166,0.206) |
| Sub. 2 | 0.003<br>(0.003,0.003) | 0.005<br>(0.004,0.005) | 0.018<br>(0.014,0.034) | 0.022<br>(0.021,0.025) | 0.154<br>(0.13,0.166) |
| Sub. 3 | 0.002<br>(0.000,0.003) | 0.006<br>(0.005,0.007) | 0.015<br>(0.012,0.026) | 0.058<br>(0.049,0.068) | 0.182<br>(0.164,0.205) |
| Sub. 4 | 0.003<br>(0.000,0.003) | 0.008<br>(0.006,0.008) | 0.043<br>(0.028,0.092) | 0.041<br>(0.024,0.051) | 0.186<br>(0.158,0.196) |
| Sub. 5 | 0.002<br>(0.000,0.003) | 0.009<br>(0.008,0.010) | 0.030<br>(0.024,0.065) | 0.032<br>(0.027,0.037) | 0.303<br>(0.271,0.323) |
| **All Sub.** | **0.003**<br>**(0.000,0.004)** | **0.007**<br>**(0.005,0.009)** | **0.022**<br>**(0.014,0.041)** | **0.034**<br>**(0.025,0.050)** | **0.198**<br>**(0.147,0.258)** |
| **$AI_0$** |  |  |  |  |  |
| Sub. 1 | 0.000<br>(0.000,0.002) | 0.001<br>(0.001,0.002) | 0.005<br>(0.003,0.007) | 0.006<br>(0.005,0.007) | 0.042<br>(0.033,0.047) |
| Sub. 2 | 0.000<br>(0.000,0.000) | 0.000<br>(0.000,0.001) | 0.005<br>(0.003,0.009) | 0.017<br>(0.013,0.019) | 0.024<br>(0.021,0.027) |
| Sub. 3 | 0.000<br>(0.000,0.001) | 0.001<br>(0.001,0.002) | 0.003<br>(0.002,0.006) | 0.009<br>(0.008,0.009) | 0.034<br>(0.028,0.040) |
| Sub. 4 | 0.000<br>(0.000,0.000) | 0.001<br>(0.001,0.003) | 0.018<br>(0.007,0.024) | 0.014<br>(0.010,0.017) | 0.037<br>(0.033,0.041) |
| Sub. 5 | 0.000<br>(0.000,0.000) | 0.001<br>(0.001,0.002) | 0.009<br>(0.005,0.014) | 0.006<br>(0.005,0.010) | 0.093<br>(0.081,0.103) |
| **All Sub.** | **0.000**<br>**(0.000,0.000)** | **0.001**<br>**(0.000,0.002)** | **0.006**<br>**(0.003,0.010)** | **0.009**<br>**(0.006,0.013)** | **0.040**<br>**(0.024,0.062)** |
| **AI** |  |  |  |  |  |
| Sub. 1 | 5.745<br>(2.504,7.369) | 14.505<br>(12.166,16.730) | 37.066<br>(33.375,41.246) | 28.311<br>(24.978,37.332) | 128.847<br>(113.873,135.39) |
| Sub. 2 | 1.124<br>(0.536,2.210) | 8.421<br>(7.398,9.537) | 44.291<br>(39.960,49.576) | 24.230<br>(21.487,39.287) | 97.084<br>(88.253,103.289) |
| Sub. 3 | 0.000<br>(0.000,2.442) | 10.437<br>(7.720,15.131) | 55.963<br>(51.028,59.763) | 27.359<br>(21.271,33.922) | 111.914<br>(102.208,121.594) |
| Sub. 4 | 0.000<br>(0.000,2.598) | 11.002<br>(9.077,13.150) | 51.713<br>(40.562,61.589) | 59.428<br>(34.030,83.187) | 117.736<br>(109.104,122.844) |
| Sub. 5 | 0.000<br>(0.000,0.544) | 12.948<br>(11.625,15.230) | 39.288<br>(34.278,45.534) | 41.695<br>(30.223,61.612) | 177.989<br>(161.927,184.464) |
| **All Sub.** | **0.000**<br>**(0.000,2.294)** | **11.396**<br>**(7.407,15.713)** | **44.566**<br>**(35.171,56.015)** | **32.03**<br>**(20.833,46.736)** | **122.077**<br>**(95.299,149.542)** |

The Activity Index (Bai et al, 2016) was designed to be robust to bias by subtracting the local mean around each axis and to measurement error by constructing the mean relative to the variability at rest. An additional calibration procedure was also introduced for ENMO to mitigate the effects of calibration bias (van Hees et al (2014)). This procedure performs a linear transformation on the raw data before computing the Euclidean norm, resulting in a calibrated version of ENMO.

### 3.7 Data labeling

Classification of physical activity types requires gold standard labels for activity, which can be obtained through direct observation or by inspection of video recordings. Obtaining gold standard labels is a labor intensive process, which often restricts the process to “in-the-lab” experiments or to a few subjects. Even gold standard labels are of different quality, depending on the accuracy and resolution at which the activity type is predicted. For example, when one is interested in predicting standing up from a chair, the duration of the activity is in the 1 to 4 second range. This makes accurate labeling even for a human observer extremely difficult. If one is interested in predicting whether the person walked or not in a particular 1 minute interval, the labels will be less accurate and will simply indicate whether the person has walked in a particular interval and approximately around what time stamp and for how long. However, even these labels can be substantially mis-aligned. This can be due to multiple factors including imperfect synchronization of clocks, time elapsed between the beginning and recording of the task, basic observer or data entry error. As an example, Figure 7 displays a portion of the acceleration data recorded at the hip for one subject. The dashed-line box is the portion of 400-meter-walk period labeled by the human observer. There is a clear 45 second shift in the label relative to the actual activity. In such cases using the original label without inspection would lead to inferior prediction algorithms and waste of resources during the modeling phase. We chose to manually inspect all “in-the-lab” data for each subject and re-label walking as periods that closely correspond to the sustained harmonic walking (SHW). The overlap between the labels provided by the human observer and the labels improved by human inspection of the data was below 80% in 18 out of 49 subjects.

**Figure 7.**
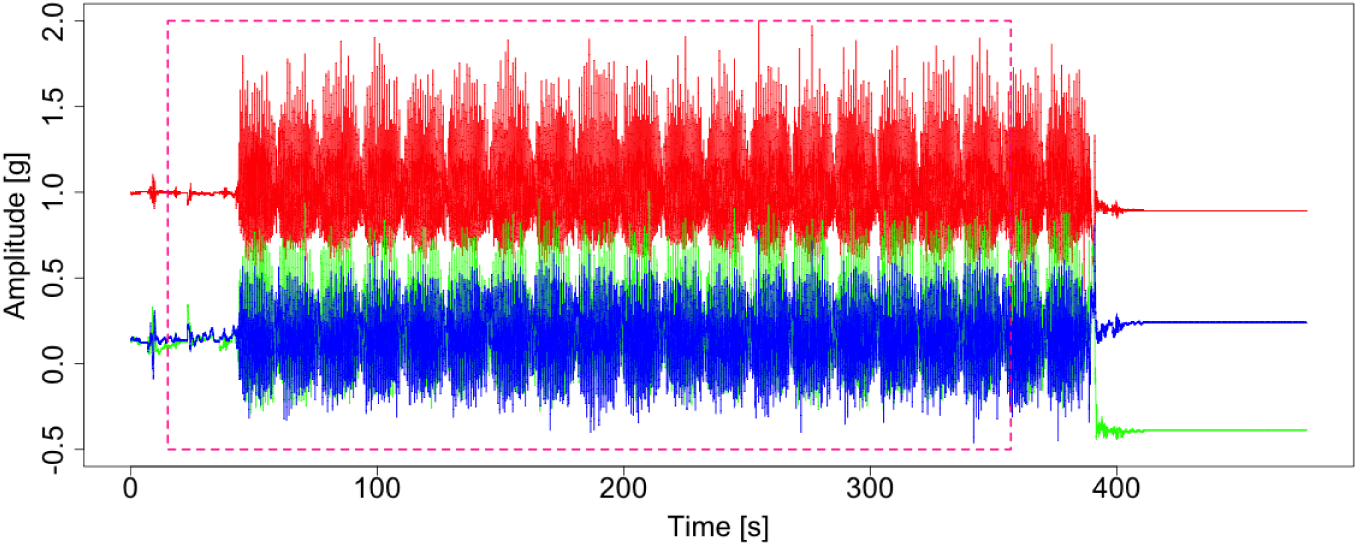
Acceleration data from three orthogonal axes at the hip collected around the time when a participant performed a 400-meter-walk activity. The dashed-line red box indicates the portion of the 400-meter-walk period identified by a human observer.

The most effective approach to proper labeling is synchronization of accelerometry data with video-recordings of the experiment (Bussmann et al, 1998). This method has been successfully used in many “in-the-lab” experiments (Godfrey et al, 2015; Del Din et al, 2016). However, using video-recordings for labeling free-living data is harder and subject to privacy considerations. Indeed, it is possible to equip participants with body-worn cameras, but video data de-identification can be quite challenging as it might contain family members, car plates, and addresses. In spite of these limitations, body-worn cameras have been used to label PA collected in the free-living environment. For example, Ellis et al (2016) used video data to train a PA classifier and Hickey et al (2017) used video data to train a walking prediction algorithm. In Hickey’s experiment, body-worn cameras were facing down to record only the feet movement.

Another, less labor intensive, approach for precise labeling of raw accelerometry data is to use landmarks introduced by the individual wearing the device. For example, in the case of wrist-worn devices, participants can clap their hands before and after each tasks. Claps result in high-amplitude, short-time spikes in the observed data that can be used to estimate timestamps for each task. For other body locations, participants can vigorously tap the device to generate proper landmarks in the data. This approach was successfully used in Straczkiewicz et al (2016). Another approach is to use the device-specific own event-markers to place labeling landmarks. Some modern PA monitors have built-in event marker buttons (e.g. Actiwatch Spectrum Plus and Pro). These event-markers generate binary indicators at the same granularity as that of the observed accelerations. When pressed, they return a value of 1 until pressed again. Initially, they were intended to mark major everyday activities (e.g. sleep) and critical events (e.g. falls), but they can be used for labeling data, as well. Unfortunately, the utility of event markers is limited by the compliance of participants (Chen et al, 2014; Boudebesse et al, 2015).

### 3.8 Synchronization of multiple PA monitors

Many studies collect data using multiple PA sensors. Synchronizing data across different monitors allows to combine information about specific human movements at multiple locations on the body (Bao and Intille, 2004; Cleland et al, 2013; He et al, 2014; Altini et al, 2015). Most devices can be set up to initialize their measurement collection at a given time (e.g. at midnight) and/or can be initialized manually. In practice, even if such approaches are used, measurements might still be desynchronized between devices. We identify two main reasons for device desynchronization.

First, most operating systems used in personal computers are not real-time operational systems. Therefore, time of execution of any command can not be precisely determined (see details in Stallings (2008)). That may result in subsecond level differences in measurements start times on multiple PA monitors. Second, the internal drift of device clocks can lead to inaccurate stamping of the time interval (Bennett et al, 2015). Such drifts are usually small (a few seconds per day) and can typically be ignored. However, when combining sub-second level data from multiple sensors in the free-living environment, the effects of the drift can have substantial side effects. To illustrate these problems, Figure 8 displays 20 seconds of data collected in the free-living environment by two monitors located on the left (top panels) and the right wrist (bottom panels). The two sensors were synchronized at the beginning of the experiment. The left and right column provide data collected on the first and seventh day of observation, respectively. The dotted vertical lines mark the end of a high-amplitude activity for each device. Although visually, data from the two devices appears to be correlated during both days, a shift in the two recordings is apparent on both days. On day 1 the time-shift is around 2.5 seconds, which is probably due to imperfect timing of device initialization. On day 7 the time shift is about 4 seconds, with the additional 1.5 seconds probably due to drift in device clocks. This drift need not be in the same direction or of the same magnitude for all devices. Such de-synchronization would have minor implications for PA summaries at the minute level collected over a 7 to 14 days period, but they can lead to substantial differences when one is interested in analyzing sub-second level data.

**Figure 8.**
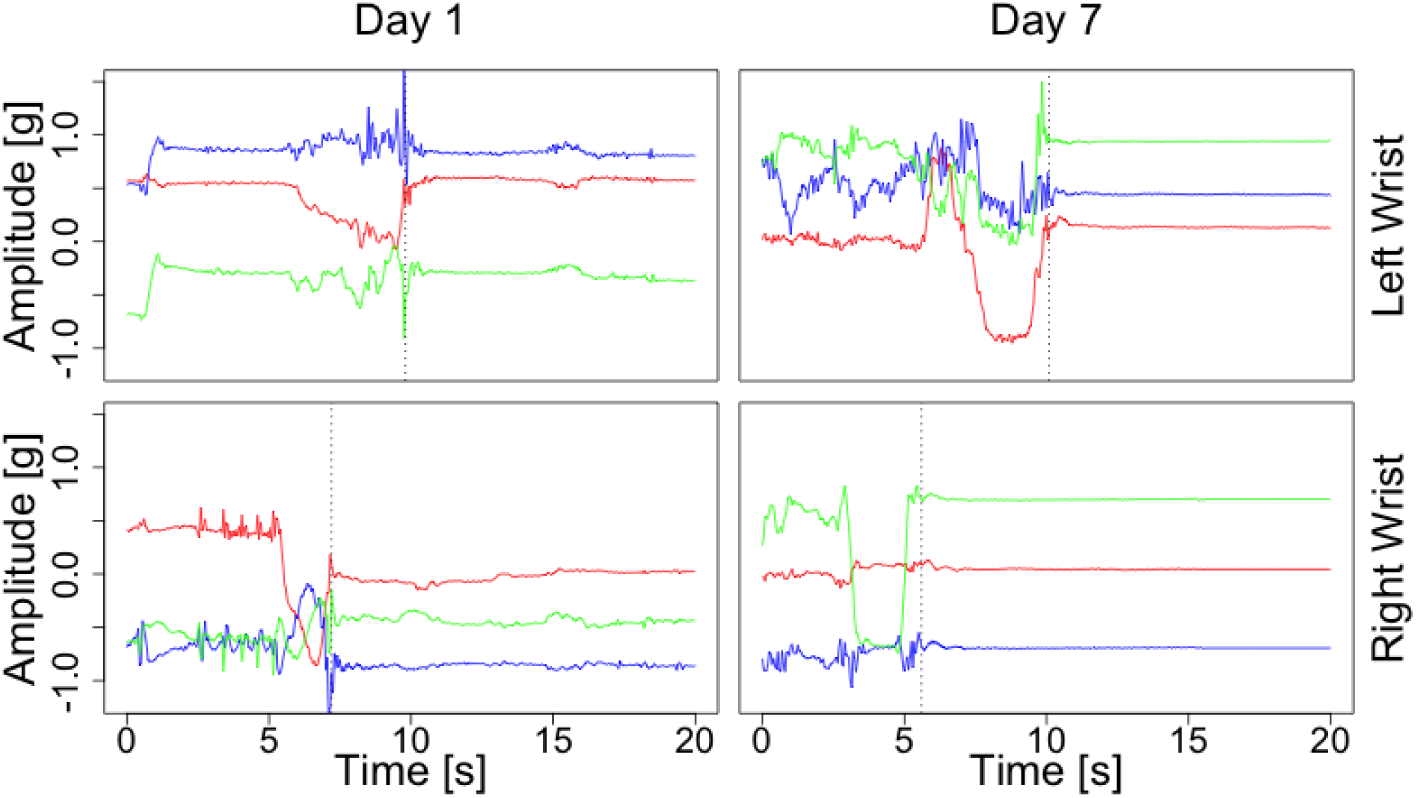
Acceleration values representing 20 seconds of data collected in the free-living environment using two monitors located on the left wrist (top panels) and the right wrist (bottom panels). The two sensors were synchronized at the beginning of the experiment. The left and right column provide data collected on the first and seventh day of observation, respectively.

The time shift introduced during device synchronization and initialization can be addressed using video recordings or landmarks. Lab experiments rarely last more than a few hours, and the effects of time drift can often be ignored. However, for data collected ‘in the free-living environment, the effects of time drift are cumulative, which can raise substantial challenges for data analysis. A possible solution could be to use devices designed specifically for parallel measurements. For example, GaitUp (GaitUp, 2017) is a system for synchronous measurement of feet movement. Alternatively, one can perform landmark-based synchronization for every time-interval in the data (e.g. every day).

## 4 Discussion

We have presented challenges related to the collection and analysis of raw, sub-second level accelerometry data. The increased granularity of observations when moving from the minute to the sub-second resolution leads to a large increase both in the volume and complexity of the data. This makes raw data harder to use than summarized data, but also holds promise of unlocking additional information glossed over by taking minute-, hour-, or day-level summaries. For example, describing gait parameters during the course of the day in the free-living environment and characterizing their potential association with health outcomes cannot be done without using sub-second level data.

Raw accelerometry data requires specialized visualization and analytic methods. This is fertile ground both for scientific researchers, as additional information is likely embedded in the raw signals, and for data scientists, as new methods and insights are becoming increasingly necessary. To start addressing this complexity we make a few points that are worth remembering: 1) activity counts are summaries of the raw data, which can depend on the device manufacturer, software version, and body location; 2) open source summary statistics are increasingly available, though more research is needed to understand their relative performance; 3) raw and summarized PA data can vary substantially with the device location, between-and within-individuals; 4) proper location choice might yield data signatures tailored to a particular study purpose; 5) device orientation can change over the course of an experiment and needs to be standardized both within- and between-individuals and devices; 6) sampling frequency can affect both raw accelerometry and summarized measurements; 7) device calibration, bias removal, and measurement error quantification can lead to higher quality data; 8) proper labeling of data is very important for training activity classifiers at the sub-second level, especially for short activities; and 9) synchronizing multiple devices must be done carefully and needs to be accounted for during the design of the experiment phase.

In spite of these challenges, the number of publications focused on raw accelerometry is continuously increasing, especially in the area of activity type classification. This is due in part to the increased popularity of these devices, their convenient design, and reduced cost. The application of raw accelerometry data in epidemiological studies is still in its infancy, though some important steps forward have been made. We anticipate that, as the interest changes to understanding the details of human movement kinematics in the free-living environment, the focus on raw data will become stronger. The number of studies that both collect and disseminate raw activity data will probably provide a huge boost to raw accelerometry data research. For example, the UK Bio-bank PA dataset is currently the largest of its kind. It was collected using the open-hardware AX3 acceleration sensor (Doherty et al, 2017).

In closing, we offer a few practical suggestions for the scientists who would like to conduct their own activity studies: 1) discuss your plans with a team that has expertise in activity research; 2) avoid the pitfalls of accelerometry research by incorporating robust, fault-tolerant designs of experiments; 3) if possible, use established protocols for data collection and pre-processing; 4) record and store the raw accelerometry data in addition to summaries, such as activity counts; 5) conduct a lab study and record the activity summaries for a well-defined group of activities in 10 to 100 individuals who are representative of the population to be studied.

## 5 Acknowledgements

The authors would like to acknowledge Annemarie Koster, PhD and Paolo Caserotti, PhD for designing the DECOS experiments.

## 6 Funding

This research was supported by Pittsburgh Claude D. Pepper Older Americans Independence Center, Research Registry, and Developmental Pilot Grant (PI: Glynn) NIH P30 AG024826 and NIH P30 AG024827. National Institute on Aging Professional Services Contract HHSN271201100605P. NIA Aging Training Grant (PI: AB Newman) T32-AG-000181. The project was supported, in part, by the Intramural Research Program of the National Institute on Aging.

**A Appendix**

